# Evaluating Representations for Gene Ontology Terms

**DOI:** 10.1101/765644

**Authors:** Dat Duong, Ankith Uppunda, Lisa Gai, Chelsea Ju, James Zhang, Muhao Chen, Eleazar Eskin, Jingyi Jessica Li, Kai-Wei Chang

## Abstract

Protein functions can be described by the Gene Ontology (GO) terms, allowing us to compare the functions of two proteins by measuring the similarity of the terms assigned to them. Recent works have applied neural network models to derive the vector representations for GO terms and compute similarity scores for these terms by comparing their vector embeddings. There are two typical ways to embed GO terms into vectors; a model can either embed the definitions of the terms or the topology of the terms in the ontology. In this paper, we design three tasks to critically evaluate the GO embeddings of two recent neural network models, and further introduce additional models for embedding GO terms, adapted from three popular neural network frameworks: Graph Convolution Network (GCN), Embeddings from Language Models (ELMo), and Bidirectional Encoder Representations from Transformers (BERT), which have not yet been explored in previous works. Task 1 studies edge cases where the GO embeddings may not provide meaningful similarity scores for GO terms. We find that all neural network based methods fail to produce high similarity scores for related terms when these terms have low Information Content values. Task 2 is a canonical task which estimates how well GO embeddings can compare functions of two orthologous genes or two interacting proteins. The best neural network methods for this task are those that embed GO terms using their definitions, and the differences among such methods are small. Task 3 evaluates how GO embeddings affect the performance of GO annotation methods, which predict whether a protein should be labeled by certain GO terms. When the annotation datasets contain many samples for each GO label, GO embeddings do not improve the classification accuracy. Machine learning GO annotation methods often remove rare GO labels from the training datasets so that the model parameters can be efficiently trained. We evaluate whether GO embeddings can improve prediction of rare labels unseen in the training datasets, and find that GO embeddings based on the BERT framework achieve the best results in this setting. We present our embedding methods and three evaluation tasks as the basis for future research on this topic.

## 1 Introduction

The Gene Ontology (GO) provides a systematic language for describing protein functions [4]. This database acts like a dictionary where each GO term has a definition explaining the biological events that it represents. Moreover, GO terms are arranged in a hierarchical tree (GO tree), where terms describing more specific biological functions are child nodes of more generic terms. A typical protein is assigned three types of GO terms: Biological Processes (BP) which identify events that involve this protein (e.g. cell division), Molecular Functions (MF) which specify the types of reactions the protein induces (e.g. kinase binding), and Cellular Components (CC) which determine the protein’s location (e.g. mitochondria) [20]. Such annotation schema let us compare the functions of two proteins by measuring the similarity of the terms assigned to them, and there have been various similarity metrics proposed for this task [26]. Traditional methods for estimating similarity scores among GO terms rely on the Information Content (IC) and the GO tree [8, 25]. The key idea is to first retrieve the shared common ancestors for two terms and their ICs, then build a function that maps these ICs to a similarity score. For example, Resnik [15] takes the maximum IC of the shared ancestors as the similarity score for two given terms.

Recently, several papers have introduced similarity metrics derived from neural network models. In principle, these neural network models solve the same problem of GO term comparison; that is, they transform related terms into comparable vectors. We will define *GO embeddings* as the vectors representing the GO terms, and *GO encoder* as the method that produces the GO embeddings. Once the GO embeddings are created, any metrics like cosine similarity or Euclidean distance can be applied to compute the similarity scores for the GO terms. From here, we can derive metrics to compare two sets of GO terms assigned to two proteins.

GO embeddings have also been used in other bioinformatics applications. Smaili <u>et al.</u> [17] used their GO embeddings to predict interactions among proteins in a Protein-Protein interaction (PPI) network. Wang <u>et al.</u> [22] jointly analyzed the PPI network and the GO embeddings to predict GO labels for unlabeled proteins in the network. Due to their wide range of applications, it is important to compare different types of GO embeddings in canonical tasks proposed by earlier works, and also study edge cases in which the embeddings may fail. In this paper, we adapt three popular frameworks in deep learning to produce additional types of GO encoders, and design three main tasks to evaluate these new encoders against the two existing encoders in [3, 17].

Typically, GO encoders are divided into two classes; they either embed the definition of a GO term or the term itself as one single entity (e.g. GO name) into vectors—we will refer to these two types as *GO definition encoders* and *GO entity encoders* respectively. For example, the definition encoder in [3] applies Bidirectional Long-short Term Memory (BiLSTM) to the GO definitions; whereas, the entity encoder Onto2vec [17] applies Word2vec [11] on axioms such as “GO:0060611 is_subclass GO:0060612" to capture relatedness of the GO names.

In this paper, we compare the methods in [3, 17] and three other neural network models: Graph Convolution Network (GCN) [5], Embeddings from Language Models (ELMo) [14] and Bidirectional Encoder Representations from Transformers (BERT) [2]. GCN, ELMo and BERT are key components in many machine learning methods, but have not yet been implemented to produce GO embeddings. GCN, ELMo and BERT have their own characteristics. GCN is an entity encoder related to Onto2vec, and applies convolutional neural network to the nodes on the GO tree. ELMo uses two BiLSTM layers, where the first layer is the input of the second layer. BERT does not apply BiLSTM or convolutional neural network, but rather models pairwise interactions for every word in the definition of a GO term.

We define three tasks to evaluate the types of GO embeddings. In Task 1, we compare the similarity score distributions for pairs of child-parent terms versus pairs of unrelated terms. This task analyzes the edge cases where the GO encoders may fail. For example, many child-parent terms located near the root node can have generic definitions which make them appear unrelated; in this case, definition encoders may not produce similar embeddings for these terms. Moreover, a high-level term can have many child nodes and so appear in many different contexts; as a result, entity encoder like Onto2vec may produce a nonspecific embedding for this term which will not closely match the embeddings of its child nodes. For all types of GO embeddings, we find that the similarity scores for child-parent terms are strongly affected in these edge cases. When two child-parent terms have low IC value, then the similarity score is not much higher than the score for two randomly chosen terms.

In Task 2, we measure the similarity scores for orthologous genes and then for interacting proteins by computing the similarity scores between the sets of terms annotating these genes and proteins. This is a canonical task evaluated in many earlier works [10]. Task 2 is a more realistic evaluation for different types of GO embeddings, because in practice, genes and proteins are not manually annotated with uninformative terms. For this reason, an encoder can perform well in Task 2 even when it does not do well in Task

1. We find that definition encoders are better than entity encoders in this task, and that performance differences among the definition encoders are small.

In Task 3, we modify the GO annotation method DeepGO [6] to take the GO embeddings as inputs, and observe which types of embeddings work best for predicting protein annotations. Surprisingly, our results show that none of the GO embeddings can increase the classification accuracy on the DeepGO dataset. In this dataset, the BP, MF and CC terms occurring below 250, 50 and 50 times are removed, so that the remaining labels are not sparse and can possibly be predicted without the additional information provided by the GO embeddings. We then consider another more difficult annotation setting to observe the performance for different types of embeddings. We design our own neural network model that uses the GO embeddings to predict labels unseen in the train data (this is zeroshot learning [18]). In this case, we find that methods with the BERT framework obtain the best accuracy. Our software is available at https://github.com/datduong/EncodeGeneOntology.

## 2 Methods

### 2.1 Training objective function and data

We use cosine similarity score to compare the embeddings for two GO terms. Embeddings for a GO term and its parent terms are expected to be comparable, and should have higher similarity scores than the embeddings for unrelated terms. Suppose a GO encoder produces the vectors *v*_*u*_ and *v*_*t*_ for the GO term *u* and *t*. We train the parameters of this encoder to max cos(*v*_*u*_, *v*_*t*_) when *u* and *t* are child-parent terms, and to min cos(*v*_*u*_, *v*_*t*_) when *u* and *t* are randomly chosen, where the cos is the cosine similarity score.

To create the training dataset, we treat the BP, MF and CC terms as one giant connected network by treating the following one-directional relationships “is a", “part of", “regulates", “negatively regulates", and “positively regulates" as the same edge type. We define a positive sample as two child-parent terms, and a negative sample as two randomly chosen terms. We select a positive sample by randomly choosing a GO term and one of its parents. We compute the Aggregated Information Content (AIC) scores [19] for these positive samples and keep only samples having scores above the median. This step ensures each remaining sample contains truly related terms. We explain AIC method in the Appendix.

We create two types of negative samples. First, we randomly select half the terms seen in the positive samples, and couple each term *c* in this set with an unrelated term *d* also seen in the positive samples. Second, we put the same term *d* with a random term *e* not found in the positive samples. This strategy helps the training process by allowing a GO encoder to observe the same terms under different scenarios [3]. We then compute the AIC scores for the negative samples, and keep samples having scores below the median. All encoders in our paper are trained on this dataset unless specified otherwise. Our dataset is available at https://github.com/datduong/EncodeGeneOntology.

### 2.2 Definition encoders

Every GO term has a definition defining the biological event that it represents; for example, the term GO:0008218 has the name *bioluminescence*, and the definition “production of light by certain enzyme-catalyzed reactions in cells". In this paper, we concatenate the name and the definition of the term into one single description. To simplify, we will refer to this concatenation as the definition of a GO term. Naturally, to represent a GO term as a vector, we can transform its definition into a vector. Below, we explain ways to do so based on the BiLSTM method in [3], and the two recent popular architectures in Natural Language Processing, Embeddings from Language Models (ELMo) [14] and Bidirectional Encoder Representations from Transformers (BERT) [2].

#### 2.2.1 BiLSTM

We describe the BiLSTM encoder in [3]. The BiLSTM network provides contextualized vectors for words in a sentence, so that the same word has different vectors depending on its positions in the sentence. Our input to BiLSTM is the word vectors produced by Word2vec. Word2vec assigns similar vectors to words with related meanings or that are likely to co-occur [11]. We use the Word2vec output in [3] which was trained on open access Pubmed papers and has word dimension in ℝ^300^.

Using Word2vec, we transform the definition of GO term into a matrix *M* where the column *M*_*j*_ is the vector for the *j*^*th*^ word. The same word is always assigned to the same vector. To capture the fact that the same word often has different meanings depending on its position in the sentence, we apply 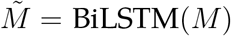. Consider the word vector *M*_*j*_ at position *j* in a sentence of length *L*. BiLSTM computes the forward and backward LSTM model to produce the output vectors 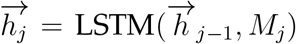 and 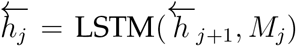 and then returns 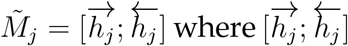 indicates the concatenation of the two vectors into one single vector.

To produce a vector representing the definition of one GO term, we take the maxpooling across the columns of 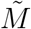, 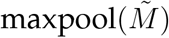 [1]. We apply a final linear transformation to this *aggregated* vector, making it into ℝ^768^ to match BERT output dimension. We train this BiLSTM model on our own dataset in section 2.1. During training, we freeze the input *M* and update only the BiLSTM parameters.

#### 2.2.2 ELMo

Embeddings from Language Models (ELMo) improves the BiLSTM encoder in two key ways [14]. First, instead of representing a whole word as a vector, ELMo represents each character in the alphabet as a vector and uses convolution filters of varying sizes to transform the alphabet vectors into a word vector. Second, ELMo trains a 2-layer BiLSTM. The first BiLSTM input are the word vectors from the character layer, and the second BiLSTM input are the output of the first BiLSTM. The final vector for one word is a weighted average of the word vector from the character layer, and the output of the first and second BiLSTM, where the weights are computed for a given specific task and thus are jointly trained with the other parameters. In this paper, we download the ELMo pretrained on Pubmed^1^, freeze the character convolution layer, and train only the two BiLSTMs. Borrowing notation from the previous section, let 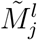 be the BiLSTM output of layer *l*. Our final vector for a word in a GO definition is 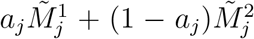 where *a*_*j*_ ∈ [0, 1] is specific to position *j* and is jointly trained with the two BiLSTMs. To encode a sentence (e.g. GO definition), we take the mean-pooling of the ELMo output matrix. ELMo is trained on the dataset in section 2.1.

#### 2.2.3 BERT

BERT is a training strategy that provides contextualized vectors for words in a sentence [2]. BERT uses the Transformer architecture [21] which models all pairwise interactions between words in a sentence. We briefly describe the key idea in the Transformer neural network model. Transformer has 12 layers, where each layer takes as input the output of the previous layer. Each layer has 12 independent units (referred to as *heads* in the original paper). We describe one Head *h* in one Layer *i*. Let *j* denote the *j*^*th*^ word in the input sentence. Consider the input “*perforation plate* is-a *cellular anatomical entity*". We use the names for these two GO terms in this example, but in the experiment we will use the complete definitions. This input is partitioned into the following tokens *[CLS] per ##fo ##ration plate [SEP] cellular an ##ato ##mic ##al entity [SEP]*, where the special token [CLS] denotes the start of the whole input and [SEP] denotes end of each sentence [2].

Each token has three types of embeddings: the token *W*, position *P* and type *T*. Token embeddings assign a token to a vector (analogous to Word2vec embeddings). Position embeddings assign a location index *j* to a vector; for example, we assign the position vector *P*_0_ to the first token [CLS]. Type embeddings assign the vector *T*_1_ and *T*_2_ to tokens in the first and second definitions respectively; for example, we assign *T*_1_ to all the tokens *[CLS] per ##fo ##ration plate [SEP]*. The input of the first layer denoted as 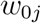 is a sum of the token, position and type embeddings corresponding to the *j*^*th*^ token. The output vector of the Head *h* in Layer *i* denoted as 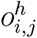 is computed as the following weighted average

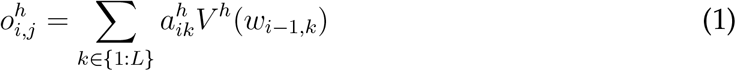

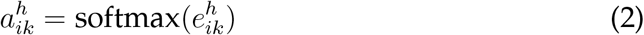

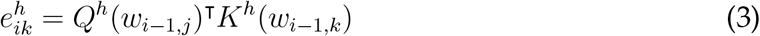

where *L* is the input length, and *V*^*h*^, *Q*^*h*^, *K*^*h*^ are transformation functions. Loosely speaking, Eq. 2 models the pairwise interaction between the *j*^*th*^ and *k*^*th*^ tokens in the sentence. To merge all the heads at Layer *i*, Transformer concatenates the output 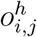 at the position *j* of all the heads, and then applies a linear transformation on this concatenated vector. The output of this linear transformation *o*_*ij*_ at position *j* is then passed onto the next Layer *i* + 1.

We now explain the two phases in BERT training strategy to train the Transformer model. In Phase 1, BERT employs a self-supervised strategy that has two key objectives. First, from the training corpus, BERT removes words from the sentences, and then predicts these removed words using the remaining words. To do this, BERT applies a classifier to the output of Layer 12 in Transformer, i.e. the vector *o*_12,*j*_. Second, BERT predicts if two sentences in a document are sequential or unrelated. Here, BERT applies a classifier to only the Layer 12 output vector *o*_12,0_ corresponding to the [CLS] token. Because Transformer models all pairwise interactions of the words, the [CLS] token can be used to represent the entire input string (e.g. definitions of two GO terms in an input string).

To train Phase 1, we create our own training corpus with respect to the context of the Gene Ontology. To create one *document*, we concatenate the definitions of all the GO terms in one single branch of the GO tree, starting from the leaf node to the root. We consider only the is-a relation, and randomly select only one parent if the node has many parents. Phase 1 will train the Transformer parameters to learn the interactions of words within the same definition, and the relationships among the definitions of terms on the same path to the root node. We use the same hyper-parameters as the original BERT, where there are 12 heads and 12 layers, and the output vector in each layer is *o*_*ij*_ ∈ ℝ^768^.

Phase 1 is often enough to produce an embedding for a sentence; for example, bert-as-service [23] takes the mean of Layer 11 output to represent the input sentence. Xiao [23] recommends the Layer 11 because they believe that values in Layer 12 are strongly affected by the two self-supervised tasks. We apply the same strategy to get the embedding for the definition of a GO term, and refer this strategy as BERT_SERVICE_. Specifically, to represent *perforation plate* as one single vector, we first retrieve Transformer Layer 11 output for the input *[CLS] per ##fo ##ration plate [SEP]*, and then compute the mean over the six tokens 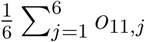. This example uses the short name of a term, but in the final model, we use the long definition.

In Phase 2 of BERT, Transformer is trained with a task-specific objective; for example, in Name Entity Recognition, the objective is to predict if a word in a sentence refers to a person, a location, or an object. In this paper, our objective is to produce a vector representing the definition of a GO term. We explore two different strategies for Phase 2.

In the first strategy, we take the mean of the Layer 12 output from the Transformer in Phase 1 to represent the definition of the GO term *c*, and then do the same for another term *p*. Denote *v*_*c*_ and *v*_*p*_ as the vector representing the definitions of term *c* and *p* respectively. We now train the Transformer on the dataset in section 2.1. The objective is to maximize the cosine similarity score, max cos(*Av*_*c*_, *Av*_*p*_) if *c* and *p* are child-parent terms, and to minimize min cos(*Av*_*c*_, *Av*_*p*_) if *c* and *p* are randomly chosen, where *A* is a linear transformation. The same Transformer in Phase 1 is used in Phase 2, so that we do not train a completely new Transformer model. We only replace the objective function, and then continue training the parameters with respect to this new objective function. We refer to this first strategy as BERT_LAYER12_. We emphasize that we train all the 12 layers of Transformer; we do not use just Layer 11 like in BERT_SERVICE_.

The second strategy is exactly the same as the first strategy, except for one key step. For a GO term *c*, we use the Layer 12 output vector *o*_12,0_ of the [CLS] token to represent its definition, and then do the same for another term *p*. Because the vector *o*_12,0_ is a function of all the words in the sentence, we reason that the [CLS] token can represent the entire input string. Abusing the notation, let us define 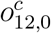 and 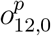 as the vectors representing *o*_12,0_ for the term *c* and *p*, respectively. We train the Transformer model to maximize 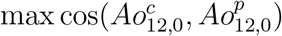 if *c* and *p* are child-parent terms, and to minimize min cos(*Av*_*c*_, *Av*_*p*_) if *c* and *p* are randomly chosen, where *A* is a linear transformation. We refer to this second strategy as BERT_CLS_. We set the final vector output for BERT_LAYER12_ and BERT_CLS_ to be in ℝ^768^, same as BERT_SERVICE_.

### 2.3 Entity encoders

Because GO terms are arranged as a hierarchical tree, we can treat a term as a single entity and encode it into a vector without using its definition. In this paper, we test Graph Convolution Network (GCN) and Onto2vec. There are other node embedding methods, but GCN has shown to work well in various applications [16, 24].

#### 2.3.1 GCN

Graph Convolution Network encodes each GO term in the tree into a vector [5]. Let *A* be the adjacency matrix, where *A*_*ij*_ = 1 if term *i* is the parent of term *j*. Compute 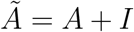, where *I* is identity matrix. Compute the degree matrix 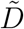, where 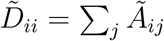. Next scale *A* into 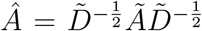. Let *W*_1_ and *W*_2_ be two transformation matrices. Define *X* to be the initial vector embedding for the GO terms, where a column in *X* corresponds to a GO vector. Before training, *X* is initialized with random numbers.

During training *X* is transformed into a new matrix *E* = *W*_2_*Â* relu(*W*_1_*ÂX*). Loosely speaking, one column *i* in *ÂX* is a summation of all its child nodes and itself, so that the information flows from child to parent under this framework [5]. *W*_1_*ÂX* then transforms this summation into a new vector. We repeat this transformation twice as in [5]. At the end, the column *i* in *E* is the vector for the *i*^*th*^ term. We set GCN to produce the final vector representation of size 768, same as BERT_SERVICE_. We train GCN using the objective function and the data in section 2.1.

#### 2.3.2 Onto2vec

Onto2vec encodes GO terms into vectors by transforming their relationships on the GO tree into sentences, which are referred to as axioms in the original paper [17]. For example, the child-parent GO terms GO:0060611 and GO:0060612 are rewritten into the following sentence “GO:0060611 is_subclass GO:0060612". Onto2vec then applies Word2vec [11] on these sentences, so that GO names occurring in the same sentence are encoded into similar vectors. Because the training *sentences* are constructed from the GO trees without GO definitions, Onto2vec can conceptually be viewed as method that encodes nodes on graph into vectors like GCN. Because the Word2vec objective function is based on cosine similarity, for Onto2vec, GO terms in close proximity will have high cosine similarity score. We set Onto2vec to produce the final vector in ℝ^768^, same as BERT_SERVICE_.

#### 2.3.3 BERT as entity encoder

Following Onto2vec, we apply BERT as an entity encoder (denoted as BERT_NAME_) where the key objective is to encode the GO names into vectors. We create training data as follows. For each GO term, we select one path from that term to the root node via only is_a relation. For each path, we split the set of GO terms into half so that they represent the *first* and *second* sentence. BERT_SERVICE_ and BERT_NAME_ use a similar idea. In BERT_SERVICE_, the training step requires GO definitions, whereas this phase in BERT_NAME_ uses only the GO names. For example, consider the path GO:0000023, GO:0005984 GO:0044262, GO:0044237, and GO:0008152. In BERT_NAME_, we format it into the input *[CLS] GO:0000023 GO:0005984 [SEP] GO:0044262 GO:0044237 GO:0008152 [SEP]*.

Next, we set the vocabularies to be learned as the GO names; that is, the token embeddings (i.e. the values to be passed into Layer 1 of Transformer) return a vector for each GO name. Then, we train the two self-supervised objectives in Phase 1 of BERT on this data, so that we can capture the relatedness among the GO names. We use the same hyper-parameters as the original BERT, where the final token embedding size is 768. After the model is trained, we treat the token embeddings as the GO embeddings. We do not take the last layer output because we do not want the contextualized vectors of the GO names which will vary depending on their locations in the input sequence and the surrounding words.

## 3 Evaluation

### 3.1 Task 1: Similarity score for two GO terms

Our GO encoders are trained so that the output vectors representing related GO terms will have high cosine similarity scores. The final GO embeddings then should produce a high cosine similarity score for any child-parent pairs. We now describe a few edge cases where it may be challenging for GO embeddings to satisfy this property.

Suppose we have two child-parent terms that describe very broad biological concepts; in this case, their definitions can appear to be very unrelated. For example, consider the term *bioluminescence* and its parent *cellular metabolic process* whose definitions are “production of light by certain enzyme-catalyzed reactions in cells" and “chemical reactions and pathways by which individual cells transform chemical substances", respectively. One definition mentions the production of light; whereas the other does not. It is then possible that the vectors representing these two terms produced by definition encoders may not have a high similarity score.

Next, consider a GO term that has many child nodes, which is often the case for terms located near root node. As an example, consider the term *cellular process* which has 59 child terms. In Onto2vec, the term *cellular process* will appear in many axioms and act like a hub, forcing all its child nodes to have very similar embeddings even when these nodes have different meanings. Subsequently, it is possible that the descendants of these dissimilar terms can also have similar embeddings. Onto2vec embeddings are then likely to return high similarity scores for unrelated terms. In GCN, our implementation builds the vector for a GO term by collecting the information from its descendant nodes. When two sibling nodes have very different descendant nodes of their own, then these two nodes can have dissimilar embeddings. Unfortunately, this strategy implies that the vector representing the parent of these two siblings cannot be very close to either of them. GCN then has the same problem observed in definition encoders.

To validate our intuition, in Task 1, we compute the cosine similarity scores between pairs of GO terms using different types of GO embeddings. We observe how the IC values of GO terms affect these similarity scores. One canonical definition for IC value of a term *g* is *IC*(*g*) = −log *p*(*g*) where *p*(*g*) is the probability that term *g* is used annotate a protein [13]. Terms having generic definitions or many child nodes tend to have low IC values because a curator is unlikely to use them to annotate proteins.

From the Human GO database, we sample 3000 child-parent pairs and 3000 pairs chosen at random. We stratify samples by the minimum IC value of the terms in a pair, and then compute the cosine similarity scores for the pairs using the embeddings of the terms. In this experiment, we compare one IC model, Aggregated Information Content (AIC) [19], against the neural network encoders (explained in the Appendix).

Figure 1 shows that AIC is in fact better than the neural network models. At each IC value interval, AIC almost always returns higher similarity scores for child-parent terms than for unrelated terms. Definition encoders and GCN achieve this result only when both child-parent terms have high IC values (Figure 1). This outcome agrees with our intuition about when the definition encoders and GCN may not properly measure the similarity scores for parent-child terms. BERT_NAME_ and Onto2vec are noticeably worse than the other methods, where the score distributions for related and unrelated pairs intersect at every IC value interval. Task 1 indicates that to generate better GO embeddings, we must integrate components of IC models into the neural network frameworks.

**Figure 1:**
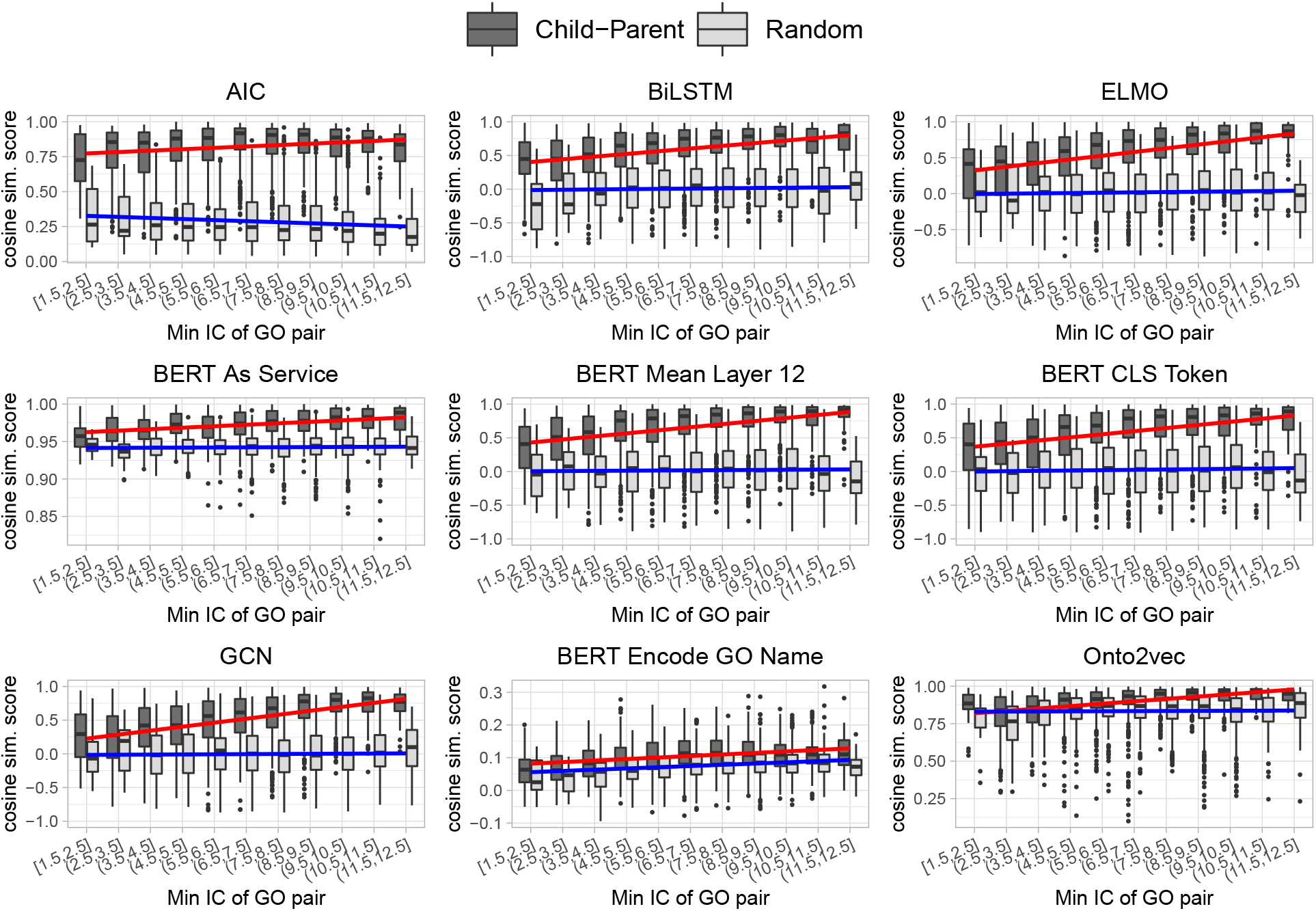
A neural network encoder’s ability to accurately classify child-parent terms is correlated to the IC values of these terms.

### 3.2 Task 2: Compare gene and protein functions

In Task 2, we compare two genes or two proteins by computing the similarity scores for the sets of GO terms annotating these genes and proteins. This is a canonical task applied in several earlier works [10]. We briefly describe the procedure to compare protein functions which is explained in more detail in [3]. We evaluate the types of GO embeddings on the orthologous gene dataset for Human–Mouse, Human–Fly, and Mouse–Fly [3], and the Human [9] and Yeast Protein-Protein Interaction (PPI) network data [12]. Because the same procedure applies to orthologous gene and PPI network datasets, we will discuss the orthologous gene dataset in more detail. The orthologous gene datasets for Human– Mouse, Human–Fly, and Mouse–Fly have 10235, 4880, and 5091 samples respectively. Each dataset contains a set of positive samples which are pairs of orthologs, and an equally-sized set of negative samples which are pairs of randomly chosen genes. Because two orthologous genes have conserved functions, their sets of GO terms are often comparable and should have a higher similarity score than two GO sets annotating two randomly chosen genes.

We apply the Best-Match Average (BMA) metric [13] to compare two sets of GO terms. This metric requires a pairwise comparison for each term in the two annotation sets. Let *A* and *B* be two sets of GO terms, and let *t*_1_ be a term in set *A*, and *t*_2_ be a term in set *B*, then the score is

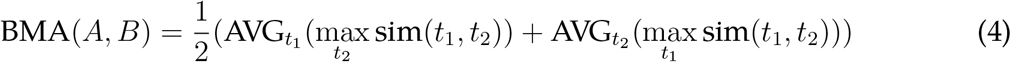

We use cosine similarity for the function sim(*t*_1_, *t*_2_). Embeddings that best estimate the similarity score for two terms will yield the most accurate BMA score. We compute the BMA score for pairs in the positive and negative samples, and then compute the Area Under the Curve (AUC) using the BMA scores as the predicted values and the true relationship between genes as the gold labels. Proper GO encoders will produce high AUC scores for each of the datasets. We repeat the same process for Human and Yeast PPI datasets, which contain 6031 and 3938 pairs respectively.

Table 1 shows the AUC for each neural network encoder. We observe that the AUCs of all the methods decrease in Human–Fly and Mouse–Fly ortholog datasets as compared to Human–Mouse dataset. We suspect that the accuracy drops because Human–Fly and Mouse–Fly datasets contain less well annotated orthologous genes. The accuracy drops more for neural network encoders than for AIC method; this result agrees with Task 1 where AIC is better at comparing GO terms with low ICs. There is no single encoder that works best for all the datasets, but the definition encoders are consistently better than the entity encoders. BERT frameworks yield the best outcome among the definition encoders. In term of model complexity and implementation, BiLSTM is the simplest definition encoder and can still perform comparably to BERT-based methods. We recommend using BiLSTM method to compare the functions of genes and proteins.

**Table 1:** AUC for classifying true orthologous genes in Human, Mouse and Fly, and interacting proteins in Human and Yeast.

|  | Orthologous gene datasets |  |  | PPI data |  |
| --- | --- | --- | --- | --- | --- |
|  | Human-Mouse | Human-Fly | Mouse-Fly | Human | Yeast |
| **Info Content** |  |  |  |  |  |
| AIC | 95.79 | **93.91** | **89.84** | 87.89 | 87.77 |
| **Defintion encoders** |  |  |  |  |  |
| BiLSTM | 95.19 | 91.49 | 80.28 | 86.61 | 88.24 |
| ELMo | 95.47 | 91.10 | 79.78 | 87.73 | 89.03 |
| BERT<sub>SERVICE</sub> | **96.72** | 92.94 | 79.62 | 88.15 | 89.00 |
| BERT<sub>LAYER12</sub> | 95.94 | 92.50 | 81.65 | **88.33** | **89.96** |
| BERT<sub>CLS</sub> | 96.05 | 90.80 | 78.99 | 86.94 | 89.27 |
| **Entity encoders** |  |  |  |  |  |
| GCN | 94.99 | 85.54 | 72.99 | 85.45 | 86.75 |
| Onto2vec | 91.80 | 82.83 | 74.41 | 79.72 | 83.98 |
| BERT<sub>NAME</sub> | 96.27 | 85.40 | 70.49 | 83.93 | 82.67 |

### 3.3 Task 3: Predict GO annotations for protein sequences

Predicting GO terms for novel protein sequences is a long-standing research problem. As of January 2020, there are 44,700 terms in the GO database; many terms are sparse, annotating only a few proteins, and cannot be readily predicted by machine learning models. For this reason, several existing statistical and deep learning annotation models are built and tested on a smaller set of GO terms that are not sparse; for example, Liu <u>et al.</u> [7] trained hierarchical classifiers on 281 and 790 labels for their yeast and human datasets, and Kulmanov <u>et al.</u> [6] trained a deep learning model on 1957 labels.

Work in other research fields have showed that the accuracy of an annotation method can improve when metadata about the labels are used as extra inputs [16, 24]. Because our GO embeddings capture the metadata about the terms such as their definitions and positions in the GO tree, we want to observe how these embeddings affect the accuracy of an existing GO annotation model. We design two sub-tasks for Task 3. First, we modify the deep learning model DeepGO [6] to take as extra inputs the GO embeddings, and evaluate our model on the same datasets in the original paper. Second, we consider a more difficult task; we build a model that uses the GO embeddings to predict labels unseen in the train datasets.

#### 3.3.1 DeepGO

In this sub-task, we integrate the GO embeddings into the GO annotation method DeepGO [6]. We use the DeepGO version which analyzes only the amino acid sequences, because we aim to observe the effect of the GO embeddings in the absence of any other factors such as PPI network data. This DeepGO version (denoted as DeepGoSeq in our paper) converts an amino acid sequence, e.g. *p* = MARS …, into a list of overlapping 3-mers MAR ARS …. Each 3-mer is assigned a vector in ℝ^128^, so that if *p* has length 1002 amino acids, then the matrix representing *p* is *E*_*p*_ ∈ ℝ^128*×*1000^. A 1D-convolution layer, 1D-maxpooling, and Flatten layer are then applied to get a vector *v*_*p*_ representing *p*, where *v*_*p*_ = flatten(maxpool (conv1d(*E*_*p*_)). To predict if the term *i* is assigned to *p*, DeepGO fits a logistic regression layer 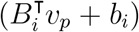, where *B*_*i*_ and *b*_*i*_ are parameters unique to label *i*. The objective loss function is binary cross entropy.

To add a type of GO embeddings into DeepGO, we make one minor change to the original model. Let *g*_*i*_ be the vector representing the term *i*, which can be retrieved by using any of our GO encoders. We concatenate *c*_*pi*_ = [*v*_*p*_ *g*_*i*_], and apply a linear transformation 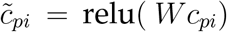 so that 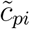 is the interaction of the sequence and the GO vector. To predict if term *i* is assigned to *p*, we fit 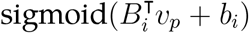 where 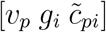 is the concatenation of the three vectors. In words, 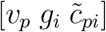 represents the information from the amino acid sequence alone, the GO metadata alone, and their interaction effects. To train the model, we keep the GO vector *g*_*i*_ constant and update only the other parameters.

We create two more baselines to critically evaluate the effect of the GO embeddings on the classification accuracy. The first baseline (denoted as +ExtraLayer) is the same as DeepGO but has one extra linear layer where the logistic regression layer now becomes 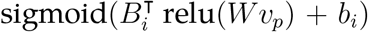. This baseline allows us to estimate the effect of adding more layers to the original model without using any GO embeddings. In the second baseline (denoted as RandomEmb), we create random embeddings where each entry is sampled from a uniform distribution *U* (−1, 1). RandomEmb acts as the control case for the actual GO embeddings, and are integrated into DeepGO in the same way that real GO embeddings are.

We train and evaluate all methods on the same DeepGO datasets. The original DeepGO paper has three datasets, one for each of the BP, MF, and CC ontology. The BP, MF, and CC train data sample sizes are 27279, 18894, and 26660 proteins, and their test data sizes are 9096, 6305, and 8886, respectively. Ancestors of the GO terms annotating a protein are added into the ground-truth label set, and then BP, MF and CC terms occurring below 250, 50, and 50 times are removed from the datasets. In total, the number of BP, MF and CC terms to be predicted are 932, 589, and 436, respectively. These BP, MF and CC labels are not sparse; the median occurrence frequencies are 365, 88, and 111 times, respectively.

In this experiment, we do not want very common labels to affect the accuracy metric; for example, the term *cell part* occurs 25,850 times and has a high AUC value 83.57 in the original DeepGO paper. We compute Micro AUC and Macro AUC, but we will focus our discussion on the Macro AUC which is the unweighted average of the per-label AUC, so that the AUC values of infrequent labels have more contributions.

We can usually obtain accurate prediction for labels that occur frequently enough in the datasets. We suspect that DeepGO may not benefit from having GO embeddings as additional inputs. Table 2 shows that this is indeed the case. Our baseline +ExtraLayer perform very well compared to DeepGO integrated with GO embeddings. Micro and Macro AUC are similar for all the types of GO embeddings, where BERT_NAME_ and Onto2vec produce competitive accuracy even when they are subpar to the other encoders in Task 1 and 2. We suspect that the parameters in DeepGO can be trained to compensate for imperfect GO embeddings; this idea is supported by the result for RandomEmb which is about the same as results for all the neural network methods. We observe that by adding only one more layer to DeepGoSeq, we can significantly increase the Macro AUC in the MF and CC data (Table 2 row 2). A more complex neural network model can probably extract more useful information form the amino acid sequences. We reserve this topic for future research work.

**Table 2:** Models are trained and tested on the same original DeepGO datasets. DeepGoSeq indicates most basic DeepGO version that analyzes only amino acid sequences of proteins. GO embeddings produced by different types of GO encoders are then integrated into the DeepGoSeq. We do not observe significant differences among the encoders, except for GCN in the BP ontology. Italicized numbers are the best baseline values, and bold numbers are the best values for the GO encoders.

|  | BP AUC |  | MF AUC |  | CC AUC |  |
| --- | --- | --- | --- | --- | --- | --- |
|  | Macro | Micro | Macro | Micro | Macro | Micro |
| **Baselines** |  |  |  |  |  |  |
| DeepGoSeq | 62.60 | 82.17 | 73.90 | 87.49 | 67.12 | 93.07 |
| +ExtraLayer | 62.82 | 82.51 | 77.39 | 88.89 | 71.03 | 93.32 |
| RandomEmb | 65.49 | 82.76 | 78.17 | 88.94 | 71.55 | 93.10 |
| **Defintion encoders** |  |  |  |  |  |  |
| BiLSTM | 64.37 | 83.16 | 76.53 | 88.83 | 70.95 | **93.56** |
| ELMo | 64.00 | 82.97 | 76.53 | 88.25 | 71.26 | 93.40 |
| BERT<sub>SERVICE</sub> | **64.95** | **83.51** | **77.74** | **89.21** | **71.85** | 93.54 |
| BERT<sub>LAYER12</sub> | 64.50 | 83.30 | 76.59 | 88.63 | 67.36 | 93.01 |
| BERT<sub>CLS</sub> | 64.49 | 83.38 | 77.08 | 88.75 | 70.45 | 93.35 |
| **Entity encoders** |  |  |  |  |  |  |
| GCN | 60.26 | 78.20 | 75.09 | 86.19 | 69.53 | 92.06 |
| Onto2vec | 64.85 | 83.41 | 76.52 | 88.90 | 69.82 | 93.31 |
| BERT<sub>NAME</sub> | 63.50 | 82.94 | 76.03 | 88.65 | 69.36 | 93.26 |

#### 3.3.2 Zeroshot experiment

When the labels to be predicted occur frequently enough, we observe that adding GO embeddings does not improve prediction accuracy. In this experiment, we consider a more difficult problem where the GO embeddings will be a key factor. We build a model that uses the GO embeddings to predict the rare labels which are not predicted in previous works [6, 7]. It is worthwhile to predict these rare labels because they are the most related to the true protein functions. For example, *perforation plate* is precise to a protein location but not its parent *cellular anatomical entity* or ancestor *cellular component*. Moreover, the GO database is frequently updated with terms being added and deleted; if we can readily predict new labels then we will not need to train the existing prediction models for each update.

Our model in this experiment relies on the zeroshot learning idea [18]. Suppose we can train a classifier *C*(*v*_*p*_, *v*_*s*_) that takes two vectors *v*_*p*_ and *v*_*s*_ representing the input *p* and the label *s* seen in the train data. Then for an unseen label *u* which can be represented as a vector *v*_*u*_, we can apply *C*(*v*_*p*_, *v*_*u*_) to classify if *p* has the unseen label *u*. Building a good zeroshot learning model is nontrivial. In this paper, our goal is not building a state-of-the-art zeroshot model; rather, we want to know which types of GO embeddings will be most suitable for future work toward this direction.

We introduce our model DeepGOZero which is based on the original DeepGO. We use the same *v*_*p*_ = flatten(maxpool (conv1d(*E*_*p*_)) to encode the amino acids. Next, we convert *v*_*p*_ into the same dimension as the GO vector *g*_*i*_ by using 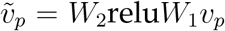. Then we create the vector *c*_*pi*_ for the final classification by concatenating four vectors 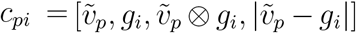. The predicted probability for label *i* is computed from three layers of fully-connected feed forward network with Relu activation. We will refer to these three layers as one layer *L*, and write the prediction as sigmoid(*L*(*c*_*pi*_)) = sigmoid(*L*(*v*_*p*_, *g*_*i*_)).

The parameters in *L* are shared for all the labels, so that terms with similar embeddings (e.g. child-parent terms) will be forced to have comparable predictions. We fix the GO embeddings as constants and train only the other model parameters. Importantly, after the model is trained, we can still predict if a protein *p* is assigned a GO term *u* which is not seen in the training label set. First, we get the vector *g*_*u*_ for this term by using any of our GO encoders. Then, we obtain *v*_*p*_ for the protein *p*, and apply the classifier layer *L* to *v*_*p*_ and *g*_*u*_. Unlike *L*, the parameters *B*_*i*_ and *b*_*i*_ in the logistic layer of DeepGO are unique for each label and cannot be applied to labels not included in the train datasets.

We now describe the datasets used to evaluate a trained DeepGOZero model. We extend the label sets in the original DeepGO data by including BP, MF, and CC terms annotating at least 50, 10, and 10 proteins, respectively (the same criteria in the original paper are 250, 50 and 50 respectively). Our BP, MF and CC datasets now have 2980, 1697 and 989 labels instead of 932, 589 and 439 labels, respectively. We keep the same number of proteins as the original datasets, so that the added labels are sparse; for example, 95% of the extra labels occur below 252, 34, and 68 times in the BP, MF and CC train data, respectively. We do not lower the selection criteria further because we want to reliably train our baseline +ExtraLayer on this entire larger datasets. Having too many sparse labels can decrease the performance of +ExtraLayer. The baseline +ExtraLayer represents the upper bound for our zeroshot learning experiment, because models trained on the complete data should be better than models trained only on part of the label sets.

We train DeepGOZero on the original DeepGO data and test on the extra 2048 BP, 1108 MF and 550 CC labels in our larger datasets. As a control case, we apply DeepGOZero with RandomEmb to estimate the accuracy due purely to chance. We will pay attention specifically to Macro AUC because it gives equal weights to rare and common labels; whereas, Micro AUC focuses more on the common labels which are often easier to classify.

In Table 3, BERT_SERVICE_ and BERT_LAYER12_ have Macro AUC closest to +ExtraLayer and furthest from RandomEmb. BERT_SERVICE_ is better than BERT_LAYER12_ for BP and MF ontology; whereas, BERT_LAYER12_ is better than BERT_SERVICE_ for CC ontology. We acknowledge that DeepGOZero is not the most effective zeroshot model to annotate protein functions; yet, BERT_SERVICE_ and BERT_LAYER12_ still come close to the upper bound +ExtraLayer. These results are proof of concept demonstrating that GO embeddings are meaningful for zeroshot learning models. We recommend the GO embeddings from BERT_SERVICE_ or BERT_LAYER12_ for future zeroshot learning methods.

**Table 3:** We add 2048 BP, 1108 MF and 550 CC labels to the original DeepGO datasets, and keep the same number of proteins. Our improved DeepGO baseline +ExtraLayer is trained and tested on the entire larger dataset, and acts as an upper bound for the other models this experiments. GO embeddings produced by different types of GO encoders are then integrated into our DeepGOZero. Models are trained on the original DeepGO datasets, and tested on the extra 2048 BP, 1108 MF and 550 CC labels in our own larger datasets. These extra labels are not seen by models during training. Italicized numbers are the upper bound, and bold numbers are the best values for the GO encoders.

|  | BP AUC | MF AUC | CC AUC |
| --- | --- | --- | --- |
|  | Macro | Macro | Macro |
| **Baselines** |  |  |  |
| +ExtraLayer | *64.64* | *72.45* | *69.31* |
| RandomEmb | 54.39 | 49.72 | 52.05 |
| **Defintion encoders** |  |  |  |
| BiLSTM | 54.91 | 61.35 | 61.45 |
| ELMo | 56.00 | 57.60 | 58.74 |
| BERT<sub>SERVICE</sub> | **62.48** | **69.96** | 62.55 |
| BERT<sub>LAYER12</sub> | 59.91 | 66.39 | **65.53** |
| BERT<sub>CLS</sub> | 55.97 | 57.78 | 58.13 |
| **Entity encoders** |  |  |  |
| GCN | 54.05 | 50.25 | 53.81 |
| Onto2vec | 56.30 | 56.82 | 52.38 |
| BERT<sub>NAME</sub> | 56.06 | 55.46 | 58.00 |

## 4 Discussion

In this paper, we design three tasks to evaluate the GO embeddings produced by the BiLSTM [3] and Onto2vec method [17]. We further introduce a few more types of GO embeddings by adapting three popular deep learning frameworks, Graph Convolution Network (GCN) [5], Embeddings from Language Models (ELMo) [14] and Bidirectional Encoder Representations from Transformers (BERT) [2].

Task 1 studies edge cases where the GO encoders may not produce accurate GO embeddings. We find that all the neural network models often fail to properly produce the embeddings for child-parent terms when these terms have low IC values (Figure 1).

In practice, manually annotated proteins are unlikely to contain terms with low IC values. To observe a realistic application of the GO encoders, in Task 2, we compare proteins by comparing the set of GO terms annotating them. We find that the definition encoders from the BiLSTM in [3], and our ELMo and BERT frameworks are better than the entity encoders BERT_NAME_, GCN, and Onto2vec in [17] (Table 1). The BiLSTM in [3] obtains competitive results against ELMo and BERT-based methods, and is easier to implement. We recommend the BiLSTM embeddings for comparing protein functions.

In Task 3, we introduce the GO embeddings into the GO annotation method DeepGO [6]. When the datasets have enough observations for each GO label, the GO embeddings do not improve the annotation accuracy. Predicting sparse labels is important for a protein, because these labels are more precise to the protein functions. We design another classifier where the GO embeddings are key factors for predicting rare labels which were excluded in the original DeepGO datasets. We train our new model on the 932 BP, 589 MF and 439 CC labels in the original DeepGO datasets, and test our model on added 2048 BP, 1108 MF and 550 CC sparse labels which were removed due to the exclusion criteria in the original DeepGO paper. BERT_SERVICE_ and BERT_LAYER12_ obtain the highest Macro AUCs and come close to the baseline trained on the entire label sets (Table 3).

Our results are proof of concept demonstrating that GO embeddings are meaningful for zeroshot learning models. Zeroshot learning has two key advantages; first, it can predict any labels in the GO database; second, zeroshot learning models are not required to be trained each time the GO database is updated with new terms. We recommend the GO embeddings from BERT_SERVICE_ and BERT_LAYER12_ for future research work on zeroshot learning.

Lastly, we emphasize that our neural network encoders are not built independently from the GO tree and the IC-methods. In BERT_SERVICE_, BERT_NAME_, GCN, and Onto2vec, the training corpus are generated from the terms that are on the same branch in the GO tree. In BiLSTM, ELMo, BERT_LAYER12_ and BERT_CLS_, we apply the AIC method to keep training samples that are very similar or are very different. For our future work, we will study other techniques to combine neural network encoders with the GO tree topology and the components of IC-models.

## 5 Appendix

### 5.1 Aggregate Information Content (AIC) Method

We describe the AIC method in Song <u>et al.</u> [19]. AIC defines a knowledge function for the GO term *t* as *k*(*t*) = 1/IC(*t*) which is used to measure its semantic value *sw*(*t*) = 1/(1 + exp(−*k*(*t*))). Here *sw*(*root*) = 1. The semantic value *sv(t)* of *t* is then defined as *sv(t)* = Σ_*p*∈path(*t*)_ *sw(p)*. Function path(*t*) contains all the ancestors of *t* and the term *t* itself. Usually, *sv*(*a*) < *sv*(*b*) when term *a* is nearer to the root than *b*. Song <u>et al.</u> [19] define their similarity score of two GO terms *a, b* as

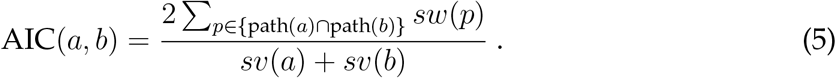

AIC(*a, b*) ranges from 0 to 1. In this model, AIC(*a, a*) = 1. When *a* and *b* only have the root node as the common ancestor, then AIC(*a, b*) = 2*/*(*sv*(*a*) + *sv*(*b*)) which depends on where *a, b* are on the GO tree.

## Footnotes

1 https://allennlp.org/elmo

